# Templated trimerization of the phage L decoration protein on capsids

**DOI:** 10.1101/2024.09.08.611893

**Authors:** Brianna M. Woodbury, Rebecca L. Newcomer, Andrei T. Alexandrescu, Carolyn M. Teschke

**Author notes:** CORRESPONDING AUTHORS: Carolyn M. Teschke, Andrei T. Alexandrescu.

## Abstract

The 134-residue phage L decoration protein (Dec) forms a capsid-stabilizing homotrimer that has an asymmetric tripod-like structure when bound to phage L capsids. The N-termini of the trimer subunits consist of spatially separated globular OB-fold domains that interact with the virions of phage L or the related phage P22. The C-termini of the trimer form a three-stranded intertwined spike structure that accounts for nearly all the interactions that stabilize the trimer. A Dec mutant with the spike residues 99-134 deleted (Dec*_1-98_*) was used to demonstrate that the stable globular OB-fold domain folds independently of the C-terminal residues. However, Dec*_1-98_* was unable to bind phage P22 virions, indicating the C-terminal spike is essential for stable capsid interaction. The full-length Dec trimer is disassembled into monomers by acidification to pH <2. These monomers retain the folded globular OB-fold domain structure, but the spike is unfolded. Increasing the pH of the Dec monomer solution to pH 6 allowed for slow trimer formation *in vitro* over the course of days. The infectious cycle of phage L is only around an hour, however, implying Dec trimer assembly *in vivo* is templated by the phage capsid. The Thermodynamic Hypothesis holds that protein folding is determined by the amino acid sequence. Dec serves as an unusual example of an oligomeric folding step that is kinetically accelerated by a viral capsid template. The capsid templating mechanism could satisfy the flexibility needed for Dec to adapt to the unusual quasi-symmetric binding site on the mature phage L capsid.

## 1. INTRODUCTION

Decoration proteins, also known as cementing proteins, non-covalently bind and stabilize the capsids of some viruses and phages, as well as serving additional functions like participating in target-cell recognition (Dedeo, Teschke, and Alexandrescu 2020). ‘Dec’ is a decoration protein encoded by phage L (Dedeo, Teschke, and Alexandrescu 2020; Gilcrease et al. 2005; Newcomer et al. 2019) that fortifies phage L virions *in vivo* against the large internal pressure resulting from genome packaging (Gilcrease et al. 2005). Phage L Dec protein can also bind to phage P22 virions, as L and P22 coat proteins and capsid structures are nearly identical (Newcomer et al. 2019; Parent et al. 2012; Tang et al. 2006). Dec has an unusual phage binding mechanism, preferring to bind as a homotrimer to specific capsid icosahedral quasi-three-fold sites that have imperfect symmetry (Gilcrease et al. 2005; Newcomer et al. 2019). Thus, the three identical protomers in the trimer must adopt slightly different conformations to optimize interactions with the capsid.

Figure 1A shows A schematic of an icosahedral capsid with the preferred quasi-three-fold sites for Dec binding that occur between hexons on the phage L capsids denoted as orange three-petal flowers. The Dec trimer has a tripod-like structure (Figure 1B) with the three globular OB-fold domains forming the legs that are the primary point of contact with the capsid (Figure 1C). The structure of the monomeric OB-fold motif was solved by solution NMR and fits well to the 4.2 Å resolution electron density map of the trimer base in the cryoEM structure (Newcomer et al. 2019). Above the tripod base, there is a C-terminal spike formed by the last ∼40 residues of each subunit with a poorer (> 6Å) resolution in the cryoEM density due to its higher mobility. The C-terminus is completely unfolded in Dec monomers indicating it only becomes structured when Dec trimerizes (Newcomer et al. 2018; Newcomer et al. 2019) (Figure 1B). Since the resolution of the spike was too low to allow cryoEM structure determination, its structure in the cryoEM data was modeled as a three-stranded β-helix structure based on its cylindrical shape and remote sequence homology to phage tail fiber proteins (Newcomer et al. 2019). Though the OB-fold domains of Dec are individually very stable, there are few if any contacts between the OB-fold domains in the capsid-bound trimer, suggesting its trimeric oligomerization state is stabilized entirely by the C-terminal spike.

**Figure 1.**
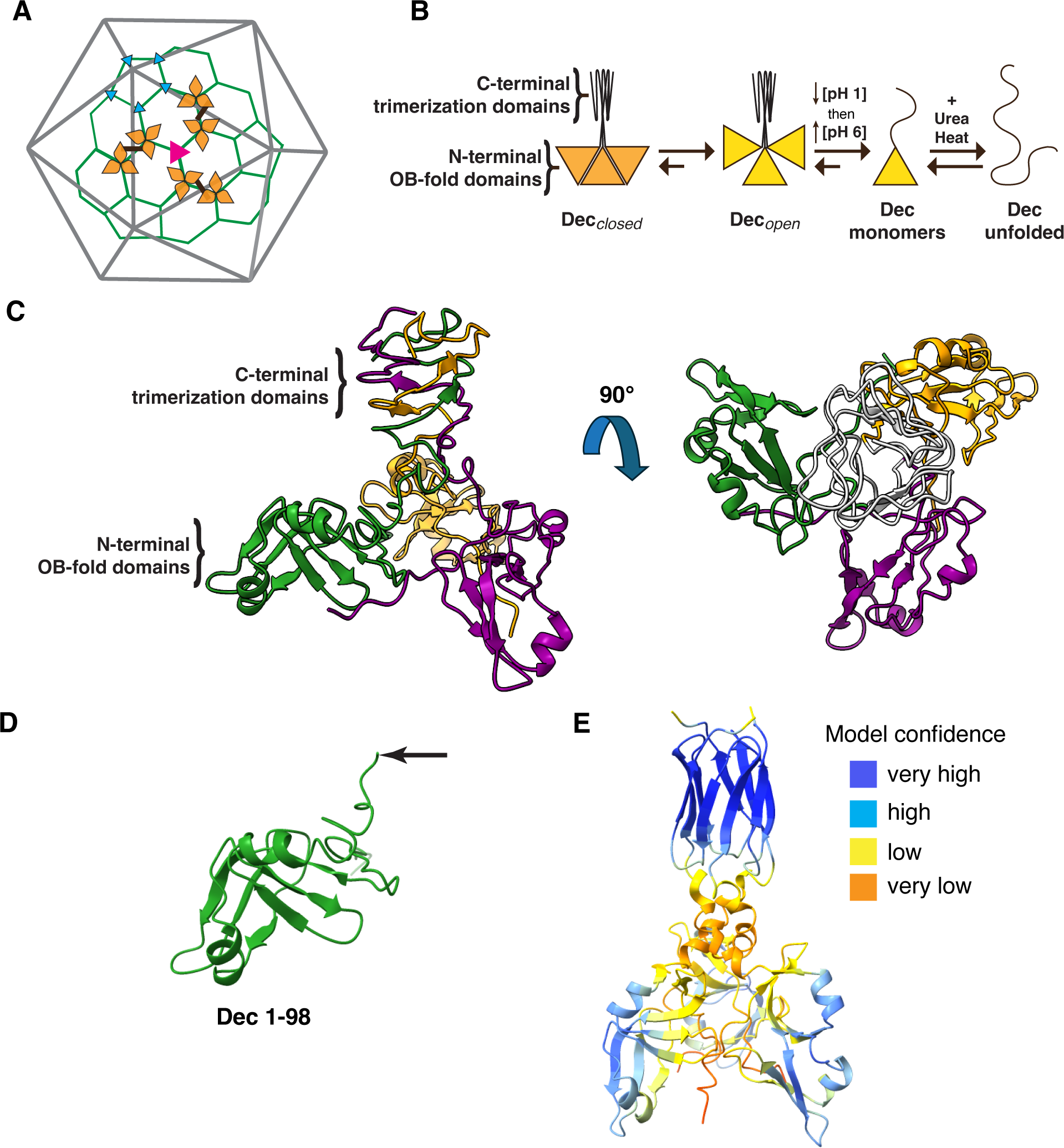
Dec structure and capsid binding sites. **A)** Diagram showing the preferred quasi-threefold binding-sites for Dec (yellow three-petal symbol) in the P22 T=7 icosahedral capsid. The icosahedral 2-fold symmetry axes are designated by black bars. The arrangement of capsid proteins into hexons and pentons is indicated by the green cages. A true 3-fold axis of symmetry is denoted by the pink triangle. Dec prefers to bind to the quasi-3 fold axes of symmetry (yellow three-petal symbol) across the 2-folds between hexons. It never binds to the quasi-3-fold symmetry axes between hexons and pentons, indicated by the small cyan triangles. **B)** Schematic illustrating the various states of the Dec protein described in this paper. In freshly purified Dec trimers (Dec*_closed_*), the OB-fold domains do not have rotational freedom. The OB-fold domains can be released from the original Dec*_closed_* conformation into the Dec*_open_* conformation by incubation at acidic pH. Further acidification unfolds the C-terminal spike and generates monomers that have a folded OB-fold domain and an unfolded C-terminal tail. When the pH is reversed to pH 6, trimerization is very slow (Figure 3B). **C)** Model of the NMR/cryoEM structure of the Dec trimer with protomers colored purple, orange, and green (Newcomer et al. 2018; Newcomer et al. 2019). In the left view, the globular OB-fold domains at the bottom of the structure contact the capsid. The top of the structure is the spike that protrudes from the capsid surface and has lower resolution in the cryoEM structure, indicative of higher flexibility. The spike was homology-modeled as a three-stranded β-helix, but its structure has not been established experimentally (Newcomer et al. 2018; Newcomer et al. 2019). An orthogonal 90°-rotated view is shown on the right, with the β-helix spike colored gray. Note that the globular OB-fold domains have no contacts with each other, and that the trimeric oligomerization state of capsid-bound Dec is maintained primarily by interactions within the spike. **D)** The Dec_1-98_ deletion mutant has the C-terminal tail genetically removed. The arrow indicates the position of residue 98. **E)** An AlphaFold2 (Jumper et al. 2021) model of the trimeric Dec structure. The spike is predicted to have anti-parallel β-barrel structure in contrast to the β-helix structure in the homology model (Figure 1C). In the AlphaFold2 model each protomer contributes 3 anti-parallel stands to the spike, with parallel β-sheet connections between the protomers. The interior of the β-barrel is not hollow but tightly packed with hydrophobic residues.

The vast majority of proteins have their native functional structure determined by their amino acid sequence according to the Thermodynamic Hypothesis (Anfinsen 1973). In rare cases, proteins can exist in multiple conformational states (Bryan and Orban 2010; Kaplan, Olson, and Alexandrescu 2021; Porter, Artsimovitch, and Ramirez-Sarmiento 2024), with the predominant species kinetically trapped in a metastable state corresponding to a sub-global energy well (Ghosh and Ranjan 2020; Baker and Agard 1994). Metastable states often occur when proteins shift between different environments (for example, a shift in pH), or undergo structural changes coupled to binding. Examples of proteins that fold into metastable states include serpin serine protease inhibitors, viral membrane fusogens such as hemagglutinin A, a number of intrinsically disordered proteins that fold only in the presence of a binding partner, and amyloidogenic proteins that often aggregate through metastable conformations (Ghosh and Ranjan 2020).

Here we investigate the folding properties of the phage L Dec trimer. Information on the folding of Dec is important both to understanding the biological functions of the protein (Gilcrease et al. 2005; Newcomer et al. 2019; Tang et al. 2006), and since the Dec trimer has become a popular carrier vehicle in nanotechnology applications such as molecular display on phage surfaces and the design of novel nanomaterials (Dedeo, Teschke, and Alexandrescu 2020; Parent et al. 2012; Schwarz et al. 2015; Goodall et al. 2021; Uchida et al. 2015). We show that Dec freshly purified from an *Escherichia coli* heterologous expression system is trimeric and that these trimers have a “closed” conformation (Dec*_closed_*) inconsistent with the cryoEM structure of the capsid-bound trimer (Newcomer et al. 2019) (Figure 1B). Dec*_closed_* trimers can be disassembled into monomers upon acidification. These Dec monomers retain a folded OB-fold domain comprised of residues 12-89 but have unfolded N- and C-termini corresponding to residues 1-11 and 90-134, respectively (Newcomer et al. 2019). In temperature melt experiments, Dec monomers and the Dec*_1-98_* C-terminal truncation mutant (Figure 1D) were completely unfolded at 63 °C in 5 M urea, with the unfolding of the OB-fold globular domain being reversible. By contrast, reassembly of Dec monomers into trimers *in vitro* required days and the trimers are consistent with a Dec*_open_* state, where the OB-fold domains have rotational freedom rather than the initial Dec*_closed_* state. We propose that *in vivo* Dec trimerization is templated by the structure of the capsid surface.

## 2. RESULTS

### 2.1 The Dec protein C-terminal tail is essential for binding to virions

In the model of trimeric Dec used for cryoEM reconstruction of the complex between Dec and phage L capsids (Newcomer et al. 2019), only the C-terminal tail forms contacts between the protomers in the trimer (Figure 1C). This predicts that in the absence of phage in solution, the globular OB-fold domains should have unrestricted rotational diffusion relative to each other (Figure 1B). The unusual oligomeric structure of Dec led us to ask how the protein assembles and the role of trimerization in binding to phage capsids.

To investigate if trimerization is essential for binding to phage capsids, a Dec truncation mutant was used to remove the C-terminal tail by insertion of a stop codon at codon 99 in the *dec* gene carried in an expression plasmid (Δ residues 99-134). (Figure 1D). The construct includes an N-terminal hexa-histidine tag, denoted as “NDec” for the full-length protein or “NDec*_1-98_*” for the truncation mutant. Note, the N-terminal His_6_-tag does not interfere with binding to capsids or trimerization (Parent et al. 2012).

We first characterized the proteins in the absence of capsids by running them on a pH 8.5 native polyacrylamide gel (Figure 2A). Freshly purified Dec (no tag) migrated on the native gel at a position consistent with being trimeric. Dec monomers (no tag) formed by incubation of Dec trimers at pH 1 followed by return to pH 6 (hereafter referred to as “Dec monomers”) (Newcomer et al. 2018) ran faster than the Dec trimers, as expected. NDec*_1-98_* ran faster than Dec monomers, at a position consistent with it being monomeric, indicating that the C-terminal tail is necessary for trimerization. Figure 2B shows ^1^H-^15^N HSQC NMR spectra of NDec monomers (black) (Newcomer et al. 2019; Newcomer et al. 2018) superimposed with the spectrum of NDec*_1-98_* (red). In an HSQC spectrum, each crosspeak is due to the amide N-H pair from a specific amino acyl residue in the protein. The close matches of chemical shifts in the superposed spectra indicate that the structure of the OB-fold is nearly identical in the monomers where the C-terminus is disordered and in NDec*_1-98_* where the C-terminus is deleted. The extra peaks in the Dec monomers compared to NDec*_1-98_* are assigned to the C-terminal tail (Newcomer et al. 2019; Newcomer et al. 2018) and have ^1^H_N_ chemical shifts between 7.8 and 8.5 ppm characteristic of unfolded structure. Thus, the C-terminal tail is not necessary for the folding of the N-terminal OB-fold domain.

**Figure 2.**
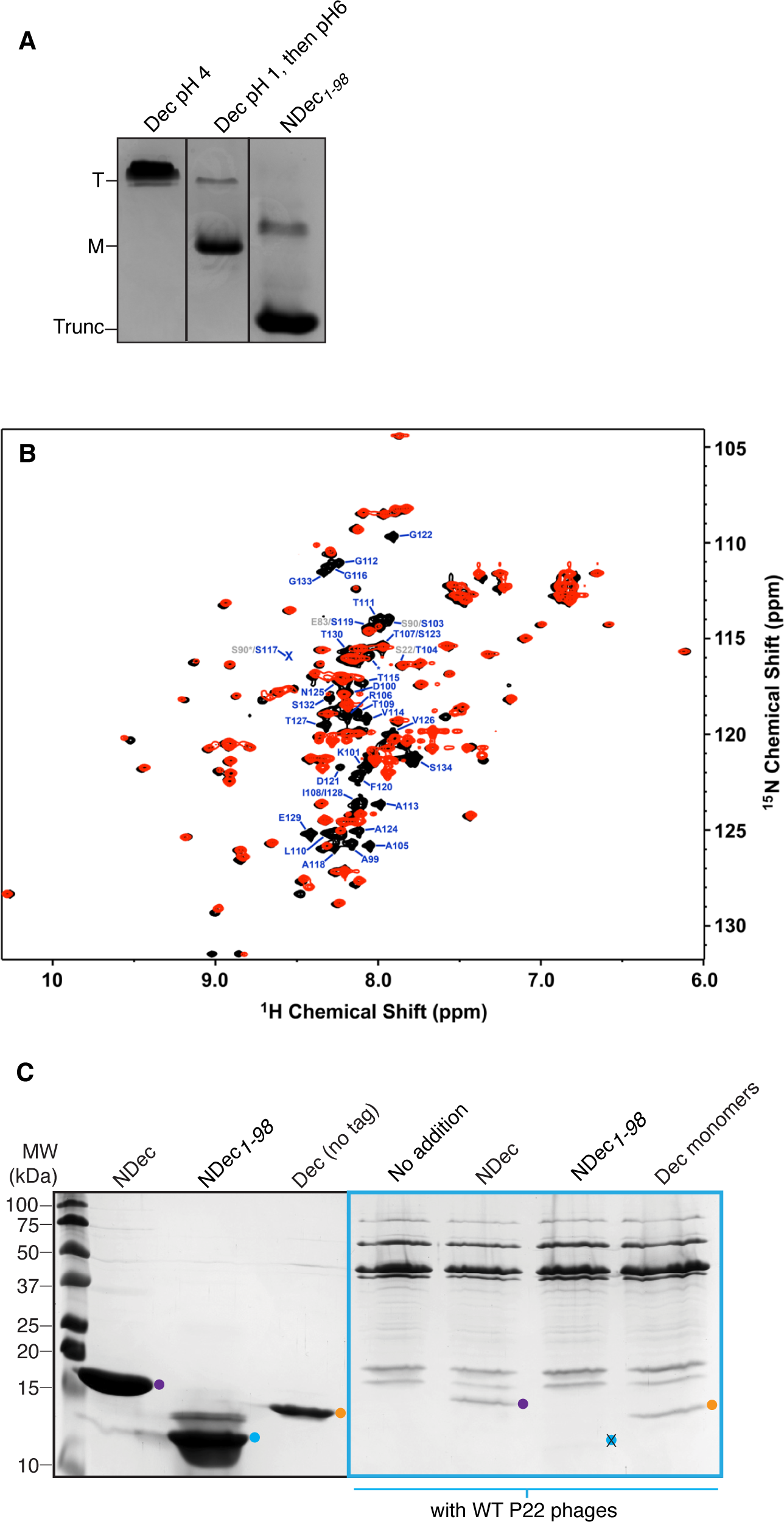
The C-terminal tail is required for capsid binding. **A)** Non-denaturing polyacrylamide gel (pH ∼8.5) showing the oligomeric state of WT Dec trimers (T), Dec monomers (M), and NDec*_1-98_* (Trunc). **B)** Superposition of NMR spectra for 1.7 mM NDec*_1-98_* (red) and 0.8 mM Dec monomers (black). Both spectra were obtained at 33 °C for samples in 50 mM NaCl, 20 mM sodium phosphate buffer, pH 6. The chemical shifts of residues in the OB-fold domains overlap indicating the structure is conserved. The extra crosspeaks in the Dec monomers are from the unfolded C-terminus (residues A99-S134), which is missing in NDec*_1-98_*. **C)** SDS-PAGE analysis of Dec proteins binding to phage P22 capsids. Purified NDec, NDec_1-98_ or Dec monomers were incubated with WT P22 phage for 30 min at room temperature then purified by CsCl step gradients. The phage band was TCA precipitated and applied to a 15% SDS-PAGE gel to assess Dec binding. Molecular weight markers (in kDa) are indicated on the left. The purified NDec, NDec*_1-98_* or Dec monomers are depicted in the left panel. The right panel, denoted by the cyan box, shows NDec, NDec*_1-98_* and WT Dec monomers that were incubated with WT P22 phages. The positions of NDec and Dec monomers on the gel are denoted by purple and orange dots, respectively. The expected position of NDec*_1-98_*, which did not bind P22, is marked by a cyan dot with an X.

To investigate if the Dec protein needs to be trimeric to bind phage virions, trimeric NDec, freshly prepared Dec monomers, and NDec*_1-98_* were each incubated with P22 phages for 15 min, after which the samples were applied to CsCl step gradients to separate free Dec from P22-bound Dec. The phage band was pulled from the gradient, the proteins were TCA precipitated, then assessed by SDS-PAGE for the presence of Dec (Figure 2C). As previously reported, the trimeric NDec bound to P22 virions (Parent et al. 2012), as did Dec monomers where the C-terminal tail is unfolded. However, monomeric NDec*_1-98_* did not bind to P22 virions, indicating that the presence of the C-terminal tail is essential for persistent binding (Figure 2C).

### 2.2 Unfolding and refolding of Dec trimers

As described above, the C-terminal tail is needed for binding of monomer Dec to phages heads, but do the Dec monomers undergo trimerization prior to capsid binding? To elucidate the mechanism of Dec-capsid binding, experiments were first performed to better understand the process of Dec folding, unfolding, and trimerization. To determine the long-term stability of WT Dec*_closed_* trimers near neutral pH, given that the trimers readily dissociate at acidic pH, the protein was incubated at pH 6 and a temperature of 33 °C. Aliquots were taken at specific times and analyzed by native PAGE. While Dec remained largely trimeric throughout the 96-hour time course, some proteolysis did occur based on the appearance of low molecular weight bands after 48 hours (Figure 3A). This result suggests that the equilibrium favors trimers at pH 6. Dropping the pH to 1 for 30 min caused trimeric Dec to dissociate into monomers (Figure 3B), indicating that the equilibrium between trimers and monomers shifts towards monomers with decreasing pH. The Dec trimers reassemble at pH 6, but over a very slow 96 h time period (Figure 3B).

**Figure 3.**
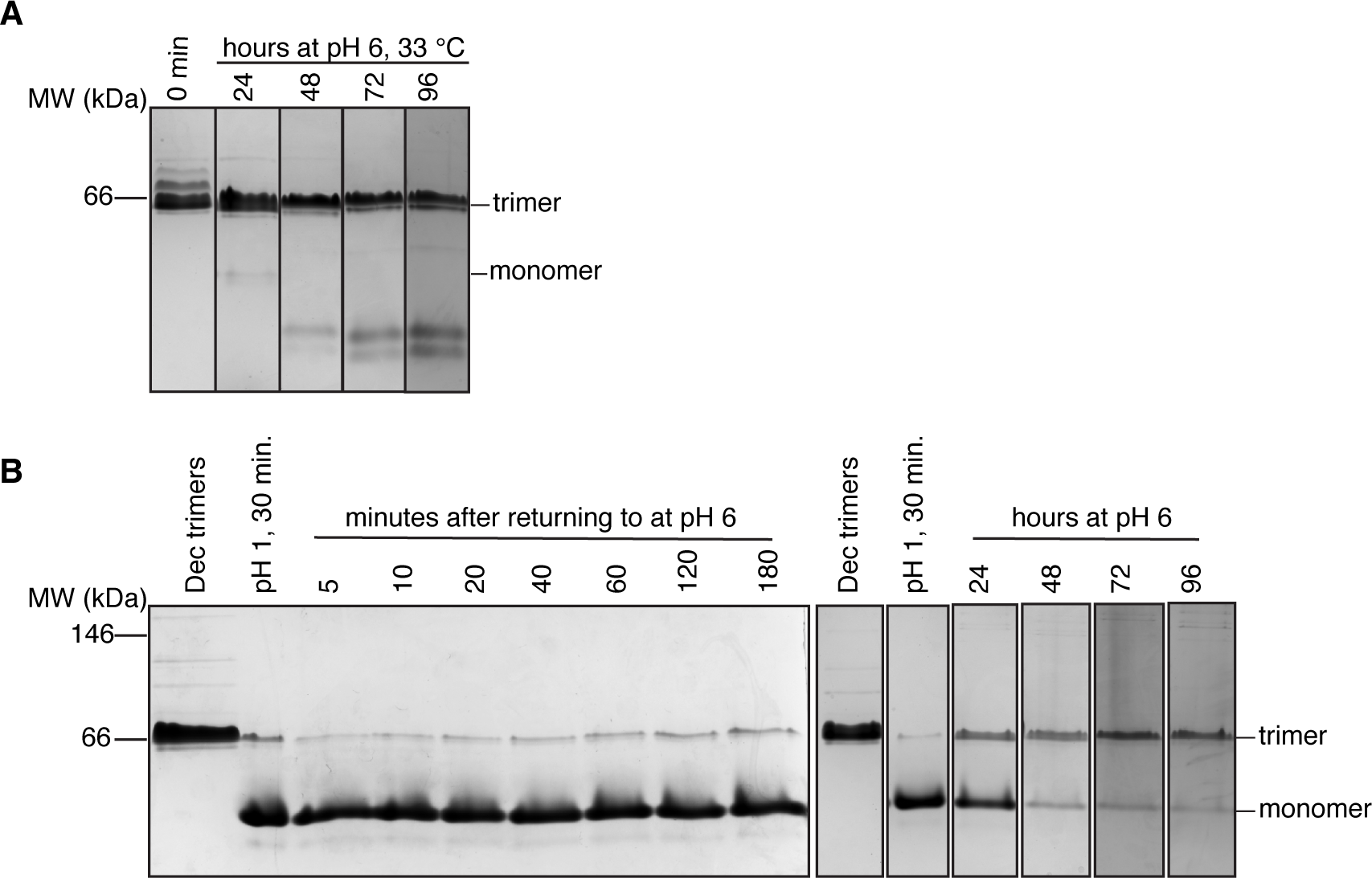
Dec trimers are very stable yet take days to assemble. **A)** WT Dec prepared in 20 mM sodium phosphate buffer pH 6 with 50 mM NaCl was thawed on ice and placed in a 33°C incubator for 96 hours. Aliquots were removed at the times indicated and analyzed by native PAGE, which showed that Dec maintains its trimeric state for up to 96 hours at pH 6 and a temperature of 33 °C. **B)** By contrast, Dec monomers dissociated at acidic pH take days to reassemble into trimers. Dec trimers (0.12 mM) were disassociated to monomers by incubation at pH 1 for 30 min. The pH was readjusted to 6, and aliquots were taken at the times indicated for analysis by native PAGE. For times up to 180 min, the aliquots were put on ice after incubation at pH 6 and run on a single native polyacrylamide gel (shown on the left). For the longer time-points, aliquots were taken from the incubation mixture every 24 hours for 4 days and run on separate non-denaturing gels. A composite of the individual gel lanes is shown on the right.

We also examined the unfolding and refolding of WT Dec using NMR ^1^H-^15^N HSQC experiments to obtain information at the residue level. At the start of the experiment, freshly purified trimeric WT Dec protein in pH 6 buffer at 33 °C showed only a few crosspeaks from the N- and C-termini (Figure 4A, top left panel). The large size of the Dec trimer (43 kDa), as well as possible additional conformational exchange contributions from a dynamic oligomerization interface, broadened most of the NMR signals in the protein. After the initial HSQC spectrum was obtained, the pH was lowered incrementally to pH 2 (indicated by the down arrow in Figure 4A), and an HSQC spectrum was collected at each increment. Obtaining each spectrum took around 75 min. Additional crosspeaks were observed as the pH was decreased, ultimately resulting in the conversion of Dec trimers into monomers at acidic pH. Of note, at pH 2 the HSQC spectrum is consistent with the protein retaining a folded OB-fold domain structure but having unfolded N- and C-termini (compare to Figure 2B). After incubation at pH 2, the pH was raised incrementally (indicated by the up arrow in Figure 4), and an HSQC spectrum was collected. With increasing pH, the spectra retained dispersed crosspeaks that were consistent with folded structure in the OB-fold domains but a spectrum similar to the initial sample with broadened NMR signals was not recovered.

**Figure 4.**
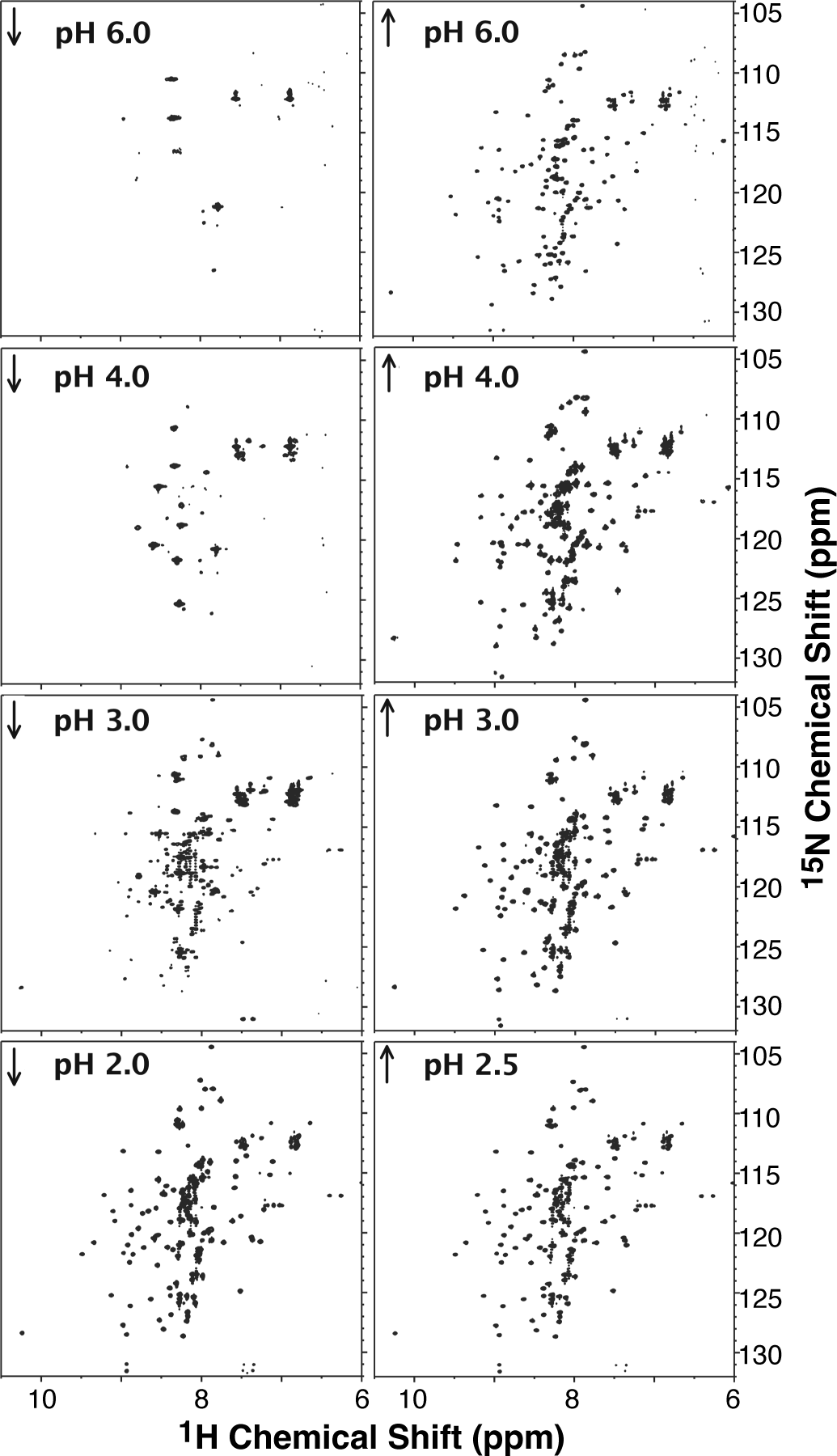
Disassembly of Dec*_closed_* trimers into Dec monomers followed by NMR. Starting with a freshly purified 0.2 mM Dec trimer sample at pH 6 and 33 °C, the pH was lowered (down arrows) and subsequently raised (up arrows) to the pHs indicated in each panel. At each pH a ^1^H-^15^N HSQC spectrum was recorded. The initial spectrum (top left) shows only a few crosspeaks indicating the OB-fold domains are interacting in the trimeric Dec*_closed_* state. As the pH is lowered, NMR signals from the OB-fold domains appear, as the trimers disassemble into monomers (see Figure 2). Adjusting the pH back to 6 does not result in the re-formation of Dec*_closed_*.

Spectra were subsequently obtained at 3 hours and 96 hours after return to pH 6 (Figure 5A, B). Even after 96 hours, the crosspeaks remained dispersed and did not return to the initial trimeric Dec*_closed_* state shown in Figure 4 (top left) where only a few NMR signals were observed. However, from the native gels shown in Figure 3B, the protein had re-trimerized by this time. The crosspeak intensities in the NMR spectrum undergo an ∼50% drop in intensity that could be due to an increase in size due to the reassociation of the monomers into trimers observed by native gels over the 96-hour time period. The chemical shifts from the C-terminus remain invariant over this time but there is a larger loss in NMR peak intensity than for the OB-fold segment (Figure 5C). This could be due to intermediate exchange broadening linked to a dynamic equilibrium between dissociated and associated forms of the C-terminal spike, and/or anisotropic tumbling of the rod-like structure. Thus, we conclude that the spectrum in Figure 5B is of Dec trimers, but in a Dec*_open_* state where the OB-fold domains have sufficient segmental rotational freedom to give NMR spectra comparable to that of the 12.9 kDa NDec*_1-98_* fragment.

**Figure 5.**
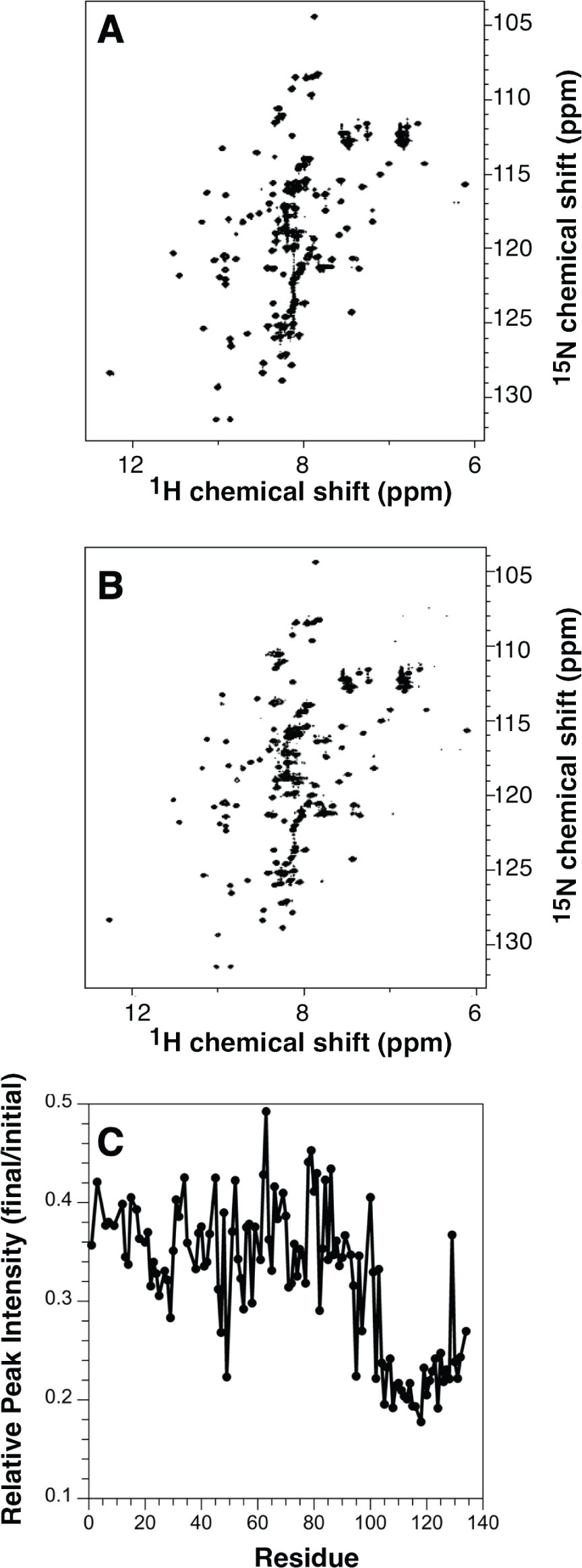
Reassembled Dec trimers have structurally independent OB-fold domains. NMR spectra of the trimeric Dec protein, dissociated at pH 2, show no changes in chemical shift positions upon raising the pH back to 6 and incubating from 3 h (**A**) to 96 h (**B**). For comparison, native PAGE show that most of the Dec monomer has reassembled to a trimer by 3-4 days, indicating that the OB-fold domains in the reassembled trimeric Dec protein undergo free rotational diffusion. We call this trimeric state Dec*_open_* to distinguish it from Dec*_colosed_*, the freshly purified trimer where NMR signals from the OB-fold segments are not observed indicating that they interact in the trimer (Figure 4, pH 6 initial) **C**) A plot of the ratio of peak intensities at 3 h compared to 96 h. Crosspeak intensities show a 2-fold decrease for residues in the C-terminal tail compared to the OB-fold domain, suggesting the C-terminal domain signals are broadened by conformational exchange or anisotropic tumbling of the spike component of the trimer.

Our data indicate that the OB-fold domain is stable even in acidic conditions of pH 1-2. Because of its high stability the OB-fold domain required incubation in 5 M urea and 63 °C to achieve thermal unfolding (Figure 6). In Figure 6A, Dec monomers were incubated in 5 M urea at 33 °C or 63 °C and compared by circular dichroism to Dec held at 33 °C without urea. Only the data from 220 to 260 nm are shown as urea absorbs strongly further into the far UV. The positive peak at 230 nm is unusual but is consistent with the presence of aromatic residues or disulfide bonds (Andersson, Carlsson, and Freskgard 2001; Haas, MacColl, and Banas 1998). As there are no cysteine residues in Dec, we attribute the peak to aromatic amino acids. In addition, there is only a single aromatic residue (F120) in the C-terminal tail, and Dec*_1-98_* also shows the positive peak at 230 nm (data not shown), indicating that the peak is due to aromatic residues in the N-terminal OB-fold domain. The peak at 230 nm decreased when Dec monomers in 5 M urea were shifted to 63 °C, consistent with Dec unfolding. When the sample was re-equilibrated at 33 °C, the protein refolded rapidly as evidenced by regaining the positive 230 nm peak.

**Figure 6.**
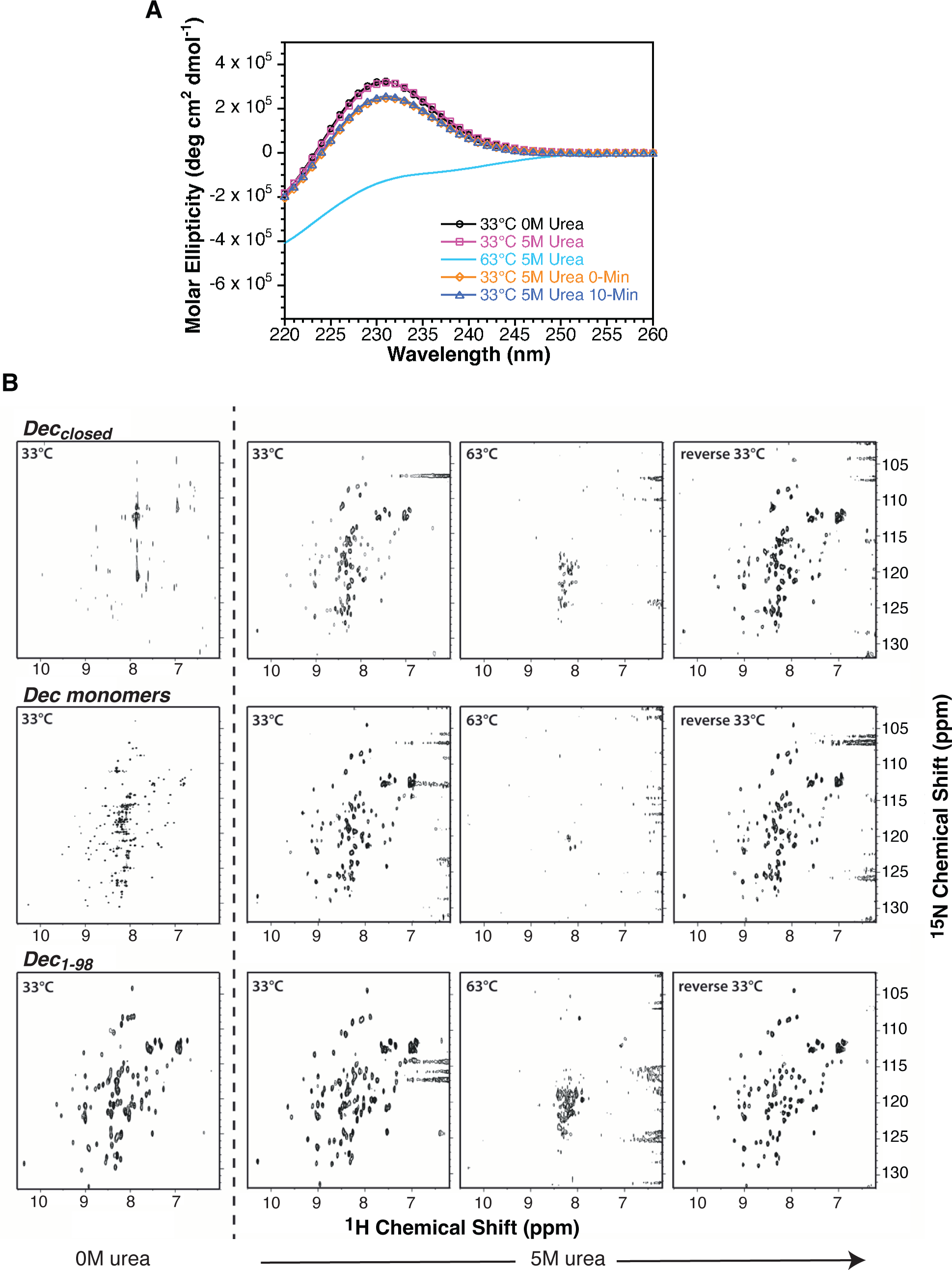
The globular OB-fold domains have a high stability to unfolding and reversible folding. **A)** Dec monomers were incubated in 0 or 5 M urea at 33 °C and circular dichroism spectra were taken. Only the data from 220-260 nm are shown due to the absorbance of urea at lower wavelengths. Addition of 5 M urea to Dec monomers does not affect the spectrum, indicating that the OB-fold domain remains folded (compare black and pink lines and symbols). 5 M urea was added for the thermal unfolding experiments because without denaturant the OB-fold domains are so stable they need higher temperatures for unfolding, which results in protein precipitation. Raising the temperature of the Dec sample in 5 M urea to 63 °C causes unfolding of the OB-fold domain (cyan line). Shifting the sample back to 33 °C allows the protein to refold the OB-fold domain, showing its unfolding is reversible (yellow and blue lines and symbols). **B)** Unfolding and refolding of Dec*_closed_* trimers, Dec monomers or Dec*_1-98_*. On the left is the spectrum of each protein in 0 M urea at 33 °C. Dec monomers and Dec*_1-98_* show the dispersed crosspeak of the OB-fold domain, while in Dec*_closed_* trimers the OB-fold domains interact resulting in few crosspeaks. In the right three panels, the proteins are in 5 M urea, starting at 33 °C. Addition of 5 M urea to Dec*_closed_* dissociates the OB-fold domains. Raising the temperature to 63 °C results in the unfolding of the OB-fold domain. Returning the samples to 33 °C leads to rapid refolding of the OB fold domain.

The high stability of the OB-fold was also observed in ^1^H-^15^N HSQC spectra (Figure 6B). The freshly purified Dec*_closed_* trimer had only a handful of resonances at 33 °C in the absence of urea. In the presence of 5 M urea there is a marked increase in resonances and the spectrum has a chemical shift dispersion typical of folded proteins. Thus, the addition of 5 M urea appears to dissociate the OB-fold domains from the Dec*_closed_* state, allowing them enough rotational diffusion freedom to be observed by NMR as is the case with acidic pH. The structure of the OB-fold domains in the presence of 5 M urea was unfolded when the temperature was raised from 33 °C to 63 °C. The crosspeaks in the ^1^H-^15^N HSQC spectrum collapse into the random coil region of the spectrum between 7.7 and 8.7 ppm and the intensities of many crosspeaks decrease due to fast hydrogen exchange from the unfolded state at high temperature. Returning the temperature back to 33 °C gave a ^1^H-^15^N HSQC spectrum indistinguishable from that at 33 °C before heating, showing that thermal unfolding of the OB-fold domains is fully reversible. The OB-fold domains of Dec monomers or Dec*_1-98_* also demonstrated reversible thermal unfolding (Figure 6B).

Taken together, these observations indicate that Dec refolding into the trimers heterologously expressed and purified from *E. coli* is not strictly reversible and that the freshly purified protein is in a state distinct from the re-trimerized protein. We are calling the initial state Dec*_closed_*, as opposed to the final trimer state Dec*_open_* obtained by reassociating acid-induced monomers at neutral pH (see Figure 1B). The operational distinction between Dec*_closed_* and Dec*_open_* is that the latter is a trimer with independently tumbling OB-fold domains as evinced by NMR signals from the majority of the residues in the protein, whereas the former Dec*_closed_* is a state where the OB-fold domains are rotationally restricted giving only a few NMR signals from the chain termini. That the NMR spectrum of the initial Dec*_closed_* is never recovered after exposure to acidic pH or high temperature indicates that this is likely a misfolded state that results from overexpression of the protein in a heterologous *E. coli* system.

### 2.3 Characterization of the Dec C-terminal tail

The structure of the Dec C-terminal tail trimers is not well-defined in the cryoEM model of Dec bound to phage L capsids (Newcomer et al. 2019). The C-terminal spike was homology modeled as a β-helix when we first reported the structure of capsid-bound Dec (Newcomer et al. 2019). We reanalyzed the Dec trimer structure using AlphaFold2 (Jumper et al. 2021). The predicted AlphaFold2 model has some striking differences from the original homology model. Rather than the C-terminus forming a parallel β-helix, the AlphaFold2 model predicts an anti-parallel β-sheet with high confidence (Figure 1E). Being agnostic about different modeling approaches, we set out to obtain experimental evidence to assess the secondary structure of the C-terminal spike using circular dichroism difference spectra. In Figure 7A, the spectrum of WT Dec trimers at pH 7.4 is compared to that of Dec monomers made by acidification (each without the His_6_-tag). Thus, the difference spectrum (trimers – monomers) should correspond to the folded trimeric C-terminal tails. Figure 7B shows the analysis of the difference spectrum using the BeSTSeL secondary structure determination suite (Micsonai et al. 2022; Micsonai et al. 2018; Micsonai et al. 2015). The fit of the difference spectrum indicates that the C-terminal tails have a high probability for anti-parallel β-sheet (Figure 7C). The fit of the difference spectrum (NDec - NDec_1-98_) also supports that the C-terminal tails have antiparallel β-sheets (data not shown). In all, the CD data are more consistent with the AlphaFold2 model than with our previous Dec model that had a parallel β-helix as the C-terminal spike. However, we note that a potential difference is that the CD data were obtained on Dec trimers freshly purified from *E. coli* that could conceivably have a different C-terminal spike structure than Dec trimers bound to capsids. Another interesting difference is that in the AlphaFold2 model the OB-fold monomers form more extensive contacts between the protomers involving loops L_23_, L_45_, and the C-terminal helix (Newcomer et al. 2019) that would probably bring the protomers too close to interact with the quasi-threefold binding site on the P22 capsid. Thus, the OB-fold protomer interactions in the AlphaFold2 model may be more relevant to the misfolded Dec*_closed_* conformation that leads to rotational restriction of the OB-fold domains than to the functional Dec*_open_* conformation where the OB-fold domains are rotationally independent. Note that the AlphaFold2 pLDDT confidence scores(Tunyasuvunakool et al. 2021) are low (orange to yellow) for the OB-fold domain segments in the Dec trimer oligomerization interface (Figure 1E).

**Figure 7.**
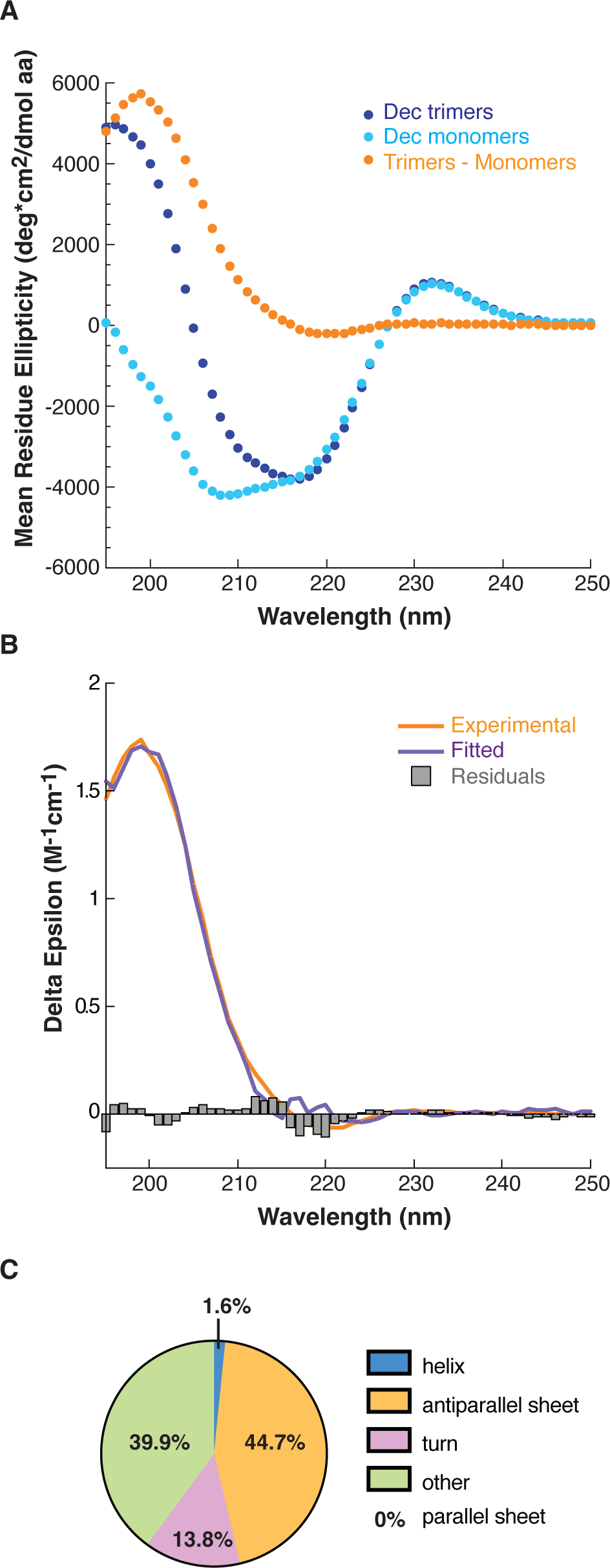
The C-terminal spike of Dec trimers is consistent with the anti-parallel β-sheet structure predicted by AlphaFold2. **A)** Circular dichroism spectra of Dec trimers (blue), Dec monomers (cyan) at 0.013 mM protomer concentration and 22 °C, and the difference spectrum of trimers-monomers (orange) corresponding to the C-terminal spike of trimers. **B)** The fit (blue line) of the difference spectrum (trimer-monomers, orange line) using BeStSel (Micsonai et al. 2022; Micsonai et al. 2018; Micsonai et al. 2015), with the residuals of the fit indicated by the gray histogram. **C)** The distribution of structure types in the fit of the difference spectrum from BeStSel.

## 3. DISCUSSION

The asymmetry of the Dec trimer structure is necessary for it to bind to coat proteins in the mature virion. The binding modality of Dec to capsids is unique. It preferentially binds with high affinity (K_d_ = 9 nM) to coat protein subunits at the capsid quasi three-fold axes of symmetry, but specifically to those between hexons that cross a two-fold axis (Figure 1A) (Newcomer et al. 2019; Tang et al. 2006; Schwarz et al. 2015; Parent et al. 2012). The quasi three-fold sites between hexons and pentons are never occupied by Dec, and the true three-fold axes can be occupied but only at a much higher Dec concentrations because of the lower binding avidity (K_d_ = 1.5 µM) at these sites (Parent et al. 2012; Schwarz et al. 2015). We suggested this unusual binding modality relates to the differing curvature of the capsid at these sites, such that the curvature of the quasi three-fold axes between hexons and pentons precludes high avidity Dec binding. Here, we hypothesize the unusual folding and assembly of Dec may be a consequence of its need to bind to an asymmetric phage capsid binding site.

### 3.1 Dec monomers are capable of binding phage capsids

The native gels in Figure 3 show that Dec trimerization is very slow, on the order of 4 days. This is an unreasonably long time for a refolding reaction, especially for a phage protein being synthesized in an infected cell that will lyse in less than an hour post-infection (Bode 1979). Our data also showed that the Dec OB-fold domain alone, using NDec*_1-98_*, is insufficient for stable binding to phage capsids (Figure 2C), suggesting that the C-terminal tail is essential for fortifying Dec in its capsid-bound state.

We tested if full-length monomeric Dec could bind to phage capsids. Dec monomers were formed by incubation at pH 1 and then the pH was shifted to 6 for 15 min. Phage P22 was added to the Dec monomer sample and incubated for an additional 30 min before being applied to the CsCl step gradient. Based on our *in vitro* refolding experiments (Figure 3B), this total incubation time (∼ 45 min) is insufficient for trimerization in solution. However, the full-length Dec monomers bound phage capsids (Figure 2C), suggesting that the C-termini must trimerize in the presence of phage capsids, thereby inducing stable binding.

### 3.2 The folding of Dec trimers is templated

The folding of the Dec protein presents several conundrums. First, the Dec OB-fold domain that directly interacts with the phage capsid is extremely stable and folds quickly. Yet the C-terminal trimeric β-sheet spike is essential for binding of Dec to the phage capsids, but its assembly into trimers is extremely slow. The timescale *in vitro* for trimer formation is several days even at the high protein (0.25 mM) concentrations used in NMR experiments, whereas the timescale of a phage L infection is less than an hour. We propose that *in vivo,* Dec monomers bind to matured phage capsids via the OB-fold domain, and if binding occurs at the proper quasi-three-fold symmetry sites between hexons, this templates the β-sheet trimer spike to lock the complex and afford it additional stability. This may also be part of the reason that Dec does not bind to immature procapsids where the particle curvature is different (Gilcrease et al. 2005). Thus, capsid-templated trimerization overcomes the slow kinetics of Dec subunit association seen in the absence of phages. It may be unfavorable for Dec subunit to associate in the absence of phage since this might lock the trimer in a conformation that is sub-optimal to bind to the unusual quasi-three-fold site on the capsid.

A second conundrum is the conformational state of trimeric Dec protein. The Dec*_closed_* conformation, which is the state of the protein when purified from cells overexpressing the protein, is not the most thermodynamically stable form of the protein. We suggest that the Dec*_closed_* conformation could be a result of the high cytoplasmic Dec concentration and the absence of phage particles. The very slow trimerization of Dec in the absence of phage capsids at the high protein concentrations during overexpression may result in misfolding to the Dec*_closed_* state. That the Dec*_closed_* state can be disrupted at acidic pH suggests that the misfolded state is stabilized by electrostatic interactions. The interactions of Dec with P22 capsids are thought to be largely driven by electrostatic interactions (Newcomer et al. 2019; Schwarz et al. 2015), so in the absence of capsid improper electrostatic interactions may be the driving force favoring the misfolded Dec*_closed_* state. Indeed, the AlphFold2 model (Figure 1E) predicts inter-protomer contacts that are not present in the Dec trimer structure bound to phage capsids (Newcomer et al. 2019). However, freshly purified Dec protein can bind to phage P22 mature virions, perhaps because interacting with capsids shifts the protein to the Dec*_open_* conformation, although we have not yet tested this hypothesis experimentally. We suggest that the rotational freedom of OB-fold domains in the Dec*_open_* trimer is functionally required to bind to quasi three-fold symmetry sites on the capsid.

Templated folding is sometimes seen in intrinsically disordered proteins (IDPs), where IDPs can be conformationally dynamic until a binding partner interacts, triggering a conformational change allowing the structure to become ordered (Toto et al. 2020). Perhaps for phage L, templated assembly overcomes the kinetic barrier for assembly of the C-terminal spike and allows for binding specificity. In terms of nanotechnology, this unique templating feature may be useful for making mixed chimeras with different proteins or probes attached to the C-termini that would only interact once bound to a capsid.

## 4. MATERIALS AND METHODS

### Cloning and protein purification

Full-length N-His_6_ Dec isotopically enriched with ^15^N was expressed and purified as previously described (Newcomer et al. 2018). To generate Dec monomers, freshly purified NDec*_closed_* trimers (pH 7.4, 14.1 mg/mL) were subjected to acidification to pH 1-2 for 30 minutes at 22 °C, and subsequent titration back to pH 6. The NDec*_1-98_* mutant was prepared using purified N-His_6_ Dec plasmid DNA from a DH5α bacterial culture that was obtained using a StrataPrep plasmid miniprep kit (Agilent Technologies). Replacing codon 99 with a stop codon was accomplished by site-directed mutagenesis, using the primer pair 5’-GTTCAGTAGCTAATGCAGAGAC and 5’-CGGGCAGTAGATAACTTATCCTAC (D’Lima and Teschke 2015; Gilcrease et al. 2005). The construct was verified through DNA sequencing by Genewiz (South Plainfield, New Jersey) and then transformed into *Escherichia coli* BL21(DE3) cells for expression.

The expression and purification of NDec*_1-98_* was similar to the purification of NDec with a few exceptions (Newcomer et al. 2018). After sedimentation, cells were resuspended in 20 mM sodium phosphate, pH 7.6, containing a 1:100 dilution of protease inhibitor cocktail (Sigma), 0.15% w/v Triton, 7 mM MgSO_4_, and 0.7 mM CaCl_2_ (lysozyme, DNase and RNase were omitted). Cells were lysed using a Constant One Shot Cell Disrupter, set to 30 psi. The purified Dec was concentrated by gentle shaking over poly(ethylene glycol) (Aldrich, average MW 20,000) in Spectra/Por dialysis tubing (8 kDa).

### NMR experiments

NMR data were collected on a 600 MHz Varian Inova spectrometer (Palo Alto, CA) equipped with a cryogenic probe, except for NMR experiments studying the combined effects of temperature and urea which were performed on a 500 MHz Bruker Avance (Billerica, MA) spectrometer. Unless otherwise noted, all samples were analyzed at a temperature of 33°C in aqueous (90% H_2_O/10% D_2_O) 20 mM sodium phosphate buffer, pH 6, 50 mM sodium chloride.

NMR experiments monitoring irreversible dissociation of Dec at acidic pH (Figure 4) were done on a 0.2 mM Dec trimer sample. NMR experiments following the reassociation of Dec from monomers to trimers with time (Figure 5) were done with a 0.25 mM sample. The sample was acidified to pH 1.9 for 30 min followed by raising the pH to 6.0 and collecting HSQC spectra at various time points, NMR experiments investigating the thermal unfolding of Dec (Figure 6B) were done on 0.2 mM, 0.6 mM and 1 mM samples of Dec*_closed_*, Dec monomers and Dec*_1-98_*, respectively. For this experiment, after collecting an initial spectrum at 33°C, urea was added to sufficiently destabilize the protein so that thermal unfolding could be observed. The 5 M urea concentration was determined using a refractometer (Carl Zeiss, Germany) according to the equation

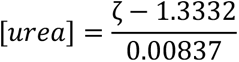

where ζ is the refractive index (Pace 1986). After addition of 5 M urea the temperature was raised to achieve unfolding and lowered to check reversibility.

NMR spectra were processed and analyzed using iNMR (http://www.inmr.net) and CCPNmr Analysis (Vranken et al. 2005). Chemical shifts of NDec*_1-_*_98_ were identified through comparison with previously published assignments of the Dec monomer (Newcomer et al. 2018).

### Purification of WT bacteriophage P22

To purify WT bacteriophage P22, *Salmonella enterica* serovar Typhimurium DB7136 (sup^0^, leu414(am), hisC527(am)) overnight cultures were diluted 1:100 in LB Super Broth [3.2% tryptone, 0.5% yeast extract, 0.5% NaCl, 7 mM NaOH] and grown to an O.D. of 0.2 at 30 °C (Botstein and Matz 1970). The cells were infected with WT P22 c1-7 (Levine 1957) (the c1-7 mutation prevents lysogeny) at a multiplicity of infection (MOI) of 0.1. The infected cultures were grown at 30°C for an additional 6 hours or until lysis occurred. The cultures were chilled on ice and chloroform was added to ensure complete cell lysis. The cell debris was pelleted by centrifugation using a Sorvall F14-6 x 250 rotor for 10 minutes at 5,000 rpm. The supernatant was transferred to an autoclaved flask containing 7% polyethylene glycol 8000 (Fisher) and 0.5 M NaCl, and gently stirred at 4 °C overnight to precipitate the phage. The phage suspension was pelleted by centrifugation at 8,000 rpm for 20 minutes at 4° C using the same rotor as above. The phage-containing pellets were resuspended overnight in dilution fluid [20 mM Tris-HCl (pH 7.6), 100 mM MgCl_2_] by gentle shaking at 4°C. The resuspended pellet was centrifuged at 10,000 rpm for 10 minutes at 4 °C using a Sorvall F18-12 x 50 rotor to remove cell debris. The resulting supernatant was ultracentrifuged at 45,000 rpm for 40 minutes at 4°C, and the phage pellet was resuspended overnight in dilution fluid by gentle shaking. Any remaining cell debris was removed via centrifugation at 10,000 rpm for 10 minutes at 4°C. The supernatant was transferred to a sterile Falcon tube and stored at 4°C.

To further purify the phage sample and to separate mature virions from procapsids, the clarified supernatant was applied to a CsCl (Sigma) step gradient (1.6 g/cc and 1.4 g/cc) with a 25% sucrose cushion (all solutions were prepared in dilution fluid). The gradients were spun in a Sorvall MX120 ultracentrifuge (Thermo Fisher) at 18 °C for 60 minutes at 30,000 rpm. The phage band was extracted from the gradient using a syringe and dialyzed against dilution fluid at 4°C overnight using a Pur-A-Lyzer Mini 6000 (Sigma) dialysis tube. The phage stock was stored at 4°C.

### Dec binding to WT phage P22

Aliquots of purified NDec*_1-98_* (pH 7.4, 1.16 mM), NDec trimers (pH 7.4, 0.98 mM) and Dec trimers (pH 4; no His_6_-tag, 0.12 mM protomers) were thawed on ice and spun in a Sorvall MX120 ultracentrifuge at 23,000 rpm for 15 minutes at 4°C to remove any aggregates. Dec monomers were prepared by incubating the Dec trimers (pH 4) sample at pH 1 for 30 minutes at 22 °C. The sample was then returned to pH 6 and allowed to incubate at 22 °C for 15 minutes (final concentration, 0.097 mM). Following this incubation period, WT P22 (final concentration of 5 x 10^12^ phage/mL) was simultaneously added to all protein samples. After mixing with the phage, the final concentration of NDec trimers was 0.069 mM protomer and NDec*_1-98_* was 0.078 mM, while the Dec monomer sample had a final concentration of 0.014 mM to decrease the possibility of trimerization. The samples were nutated at 22 °C for 30 minutes then applied to CsCl step gradients, as detailed above. The phage bands were puncture harvested using a syringe and dialyzed overnight, as above. The proteins were TCA precipitated, resuspended in 3X SDS sample buffer and analyzed on a 15% SDS polyacrylamide gel to determine if Dec bound to the WT P22 virions.

### Time course of Dec trimerization and Dec stability

An aliquot of Dec trimers (pH 4, 0.12 mM initial concentration of protomers), in which the N-terminal His_6_-tag has been cleaved, was thawed on ice. Acid-induced unfolding of the Dec trimers was achieved by adding small µL aliquots of 1N HCl followed by vortexing until the pH decreased to 1. The pH 1 sample was incubated at 22 °C for 30 minutes, followed by returning to neutral pH by slowly titrating in 1N NaOH until the pH reached 6, where the final protein concentration was 0.093 mM. The pH 6 sample was incubated at 33 °C and small aliquots were taken at specified time points for analysis by native PAGE. The native PAGE gels were prepared in-house using a 4.3% stacking gel and 15% separating gel. The native PAGE gels were run at 300V and 10 mA for 3 hours at 4°C to allow for sufficient separation of Dec trimers from Dec monomers.

To assess the long-term stability of Dec trimers, WT Dec (pH 6, 0.21 mM) without an N-terminal His_6_-tag was incubated at 33°C for 96 hours. A small aliquot was removed at specified time points and assessed by native PAGE to determine the extent of protein denaturation that occurs with time.

### Circular Dichroism of Dec

Purified Dec trimers (pH 4) and Dec monomers, prepared as above, were diluted in distilled water to a final protomer concentration of 0.2 mg/ml (0.013 mM). CD spectra were acquired using a 0.1 cm path length quartz cuvette (Starna Cells, Inc.) on a Chirascan V100 spectrometer (Applied Photophysics). The CD spectra were collected from 190 to 260 nm at 22 °C with a bandwidth of 3 nm, 1 nm intervals and a time-per-point averaging of 5 seconds. The oligomeric state of the WT Dec and WT Dec monomers was verified by native PAGE to ensure that the CD spectra correspond to trimers and monomers, respectively.

## ACKNOWLEDGMENTS

We thank Helen Belato for her help with the NMR Dec reassembly experiments. This work was supported by NIH grant R01 GM07661 to CMT and by a grant from the UConn Research Foundation to ATA and CMT.

## AUTHOR CONTRIBUTIONS

A.T.A and C.M.T conceived of the project, directed the work, analyzed data and wrote the manuscript. B.M.W. and R.L.N. did the experiments, analyzed the data, and contributed to the writing of the manuscript.

## ABBREVIATIONS

cryoEM: cryogenic electron microscopy
WT: wild type
Dec: phage L decoration protein
Dec*_closed_*: trimeric form with the three OB-fold domains interacting
Dec*_open_*: trimeric form with the three OB-fold domains not interacting
NDec: Dec construct with an N-terminal His_6_ tag
Dec monomers: Dec trimers incubated at pH <2 and returned to pH 6
OB-fold: oligonucleotide/oligosaccharide-binding fold
NMR: nuclear magnetic resonance
HSQC: heteronuclear single-quantum coherence
SDS: sodium dodecyl sulfate;
TCA: trichloroacetic acid.

